# Guide assignment in single-cell CRISPR screens using crispat

**DOI:** 10.1101/2024.05.06.592692

**Authors:** Jana M. Braunger, Britta Velten

## Abstract

**Motivation:** Pooled single cell CRISPR screens have emerged as a powerful tool in functional genomics to probe the effect of genetic interventions at scale. A crucial step in the analysis of the resulting data is the assignment of cells to gRNAs corresponding to a specific genetic intervention. However, this step is challenging due to a lack of systematic benchmarks and accessible software to apply and compare different guide assignment strategies. To address this, we here propose crispat (CRISPR guide assignment tool), a Python package to facilitate the choice of a suitable guide assignment strategy for single cell CRISPR screens.

**Results:** We demonstrate the package on four single cell CRISPR interference screens at low multiplicity of infection from two studies, where crispat identifies strong differences in the number of assigned cells, downregulation of the target genes and number of discoveries across different guide assignment strategies, highlighting the need for a suitable guide assignment strategy to obtain optimal power in single cell CRISPR screens.

**Availability and Implementation:** crispat is implemented in python and the source code, installation instructions and tutorials can be found at https://github.com/velten-group/crispat. Code to reproduce all findings in this paper is available at https://github.com/velten-group/crispat_analysis.

**Contact:** Britta Velten

**Supplementary Information:** Supplementary Information can be found in the Supplementary Information file.

## Introduction

Pooled single-cell CRISPR screens are a powerful tool for systematically gaining new insights into the functional consequences of genetic perturbations in high-throughput analyses. In such screens guide RNAs (gRNAs) are introduced into cells expressing Cas proteins and can direct the Cas proteins to a target site in the DNA that can be altered, suppressed or activated^1^. Single-cell sequencing then allows direct read-outs of molecular changes induced in each cell as well as markers linked to the gRNAs that are present in the cell, e.g. using barcodes or direct guide capture^2,3^.

To extract biological insights about gene regulatory mechanisms from single-cell CRISPR screens a first step in the computational analysis of the resulting data is guide assignment. Hereby, cells are assigned to specific gRNAs and corresponding genetic perturbations based on gRNA counts in each cell. This step can be challenging due to background contamination and sparsity of single cell measurements and the confidence of the assignment typically varies across cells. Suitable thresholds on the measured gRNA counts per cell have to be found to distinguish background noise from actual guide presence. Currently, a wide range of different guide assignment strategies and thresholds are used^4–13^ without guidance on what strategy to choose when. A systematic comparison of multiple strategies is hindered due to the lack of easily accessible software that could facilitate the application and comparison of different guide assignment strategies.

Here, we address this by developing crispat (CRISPR guide Assignment Tool), a well-documented Python package that implements 11 different guide assignment methods and facilitates their comparison. The package includes commonly used methods as well as 5 novel approaches with a focus on low multiplicity of infection (MOI) CRISPR screens, where a majority of transduced cells contain a single gRNA. We compared the performance of the methods in CRISPR interference (CRISPRi) screens with single-cell gene expression read-outs based on the number of uniquely assigned cells, the target gene downregulation in assigned cells, as well as the number of total and false discoveries.

## Methods

The starting point for the python package crispat are count matrices of the gRNA measurements in each cell obtained after aligning the sequenced gene and gRNA reads, e.g. using 10X Genomics Cell Ranger or kallisto^14^. Different file formats are supported by crispat, including file formats generated by the widely used Cell Ranger platform or a csv file containing the guide counts for every gRNA and cell. Based on this data, crispat assigns every cell to one or multiple gRNAs, providing users with a choice of 11 different guide assignment methods. These consist of literature-based methods as well as extensions thereof and can be grouped into 4 categories based on the information which is used for the assignment (Fig. 1). The first group of methods (*independent*) relies only on the individual measurement for each gRNA and cell, by checking whether an entry is higher than a user-defined threshold. While this is a simple strategy, it is widely used in applications with different thresholds^4,7–9^.

**Figure 1:**
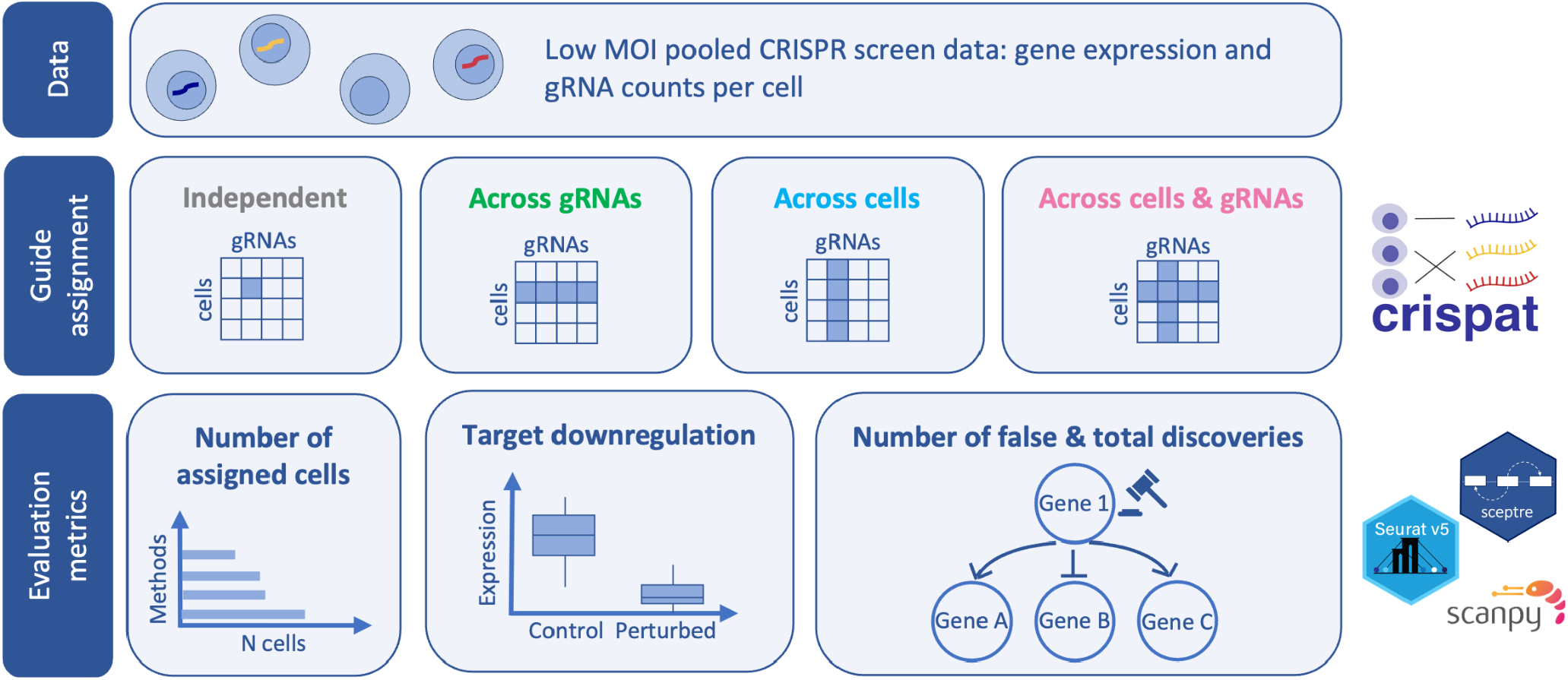
Overview of the guide assignment using crispat and the metrics considered for the comparison of different methods.

A second group of methods (*across gRNAs*), considers information for all gRNAs in a cell, e.g. by using the relative frequency of the most abundant gRNA in comparison to the total number of counts from all gRNAs in a cell for the assignment^6,10,11^. Similarly to the first group of methods, a suitable threshold needs to be found to determine when the most abundant gRNA in a cell is considered to have a substantially higher relative frequency compared to the other gRNAs in that cell and is assigned to the cell. A third group of methods (*across cells*) determines a suitable threshold for every individual gRNA by comparing its count values across all cells, e.g. using a two-component mixture model to distinguish cells with background counts from transduced cells containing the gRNA. Examples include a Gaussian-Gaussian mixture model employed by 10X Genomics Cell Ranger and a Poisson-Gaussian mixture model proposed by Replogle et al.^5^. A last group of methods (*across cells and gRNAs*) in a similar manner uses the heterogeneity across cells, e.g. to determine suitable thresholds based on mixture models per gRNA, but in addition takes into account the total amount of gRNA counts per cell. An example for this is the Poisson model accounting for the total guide counts in a cell as library size in a generalized linear model^13^. Other methods in this group directly operate on the relative frequencies of the most abundant gRNA in the cell, e.g., the “2-Beta” and “3-Beta” mixture models replace the user-defined threshold on the relative frequencies of the most abundant gRNA per cell by a learnt threshold per batch. An overview of all 11 assignment methods is shown in Supplementary Table 1 and details on every assignment method can be found in the Supplementary Information and the crispat documentation. While some methods are specifically designed for low MOI CRISPR screens, others can also be employed for screens conducted at a high MOI (Supp. Table 1). All methods relying on mixture models were implemented using pyro (version 1.8.6)^15^ and mixture models based on previous approaches were compared against their original implementations to ensure that they yield similar results (Supp. Fig. 1).

To compare different guide assignment methods using crispat we focused on four criteria: (i) number of cells with single gRNA assigned, (ii) target gene downregulation in the assigned cells, and (iii) number of discoveries defined as differentially expressed genes in cells assigned to a specific perturbation compared to cells assigned to a non-targeting gRNA (control cells), as well as (iv) number of false discoveries defined as differential expression changes between cells assigned to one non-targeting gRNA vs. all other control cells (Fig. 1). To this end, crispat can be used to compare the number of assigned cells across methods as well as the overall agreement of all assignments between different methods. In addition, the assignments can serve as direct input to differential expression tests, e.g. as demonstrated in the tutorials for Seurat^16^, scanpy^17^ and the R package SCEPTRE^18^ (Fig. 1).

## Results

To demonstrate the use of crispat, we applied it to a large-scale single-cell CRISPR interference screen, where 686,464 K562 cells were transduced with gRNAs targeting 2,285 different genes^5^. This enabled us to conduct a systematic comparison of the 11 methods implemented in crispat for guide assignment and evaluate the impact of the method choice on the power of the screen. For methods that require a user-defined threshold (Supp. Table 1) we additionally tested and compared a range of different thresholds and selected the threshold giving the highest number of significant target genes for each method (Supp. Fig. 2).

For the following analysis, we focus on uniquely assigned cells defined as cells for which exactly one gRNA was assigned as is common practice in low MOI screens^5^. While some methods such as e.g. the maximum assignment are designed for low MOI screens and assign at most one gRNA per cell, other methods such as the assignment based on a Gaussian-Gaussian mixture model resulted in many cells with more than one gRNA assigned (Fig. 2A). The number of uniquely assigned cells varied strongly from less than 150,000 to more than 600,000 cells (Fig. 2A) with 5,000 to 20,000 cells assigned to non-targeting control gRNAs (control cells) and a median of 100-200 cells per targeting gRNA (Supp. Fig. 3). Overall, we observed a high consistency in the guide assignments across methods with lower Jaccard indices mostly due to differences in the number of cells with single gRNA assigned (Fig. 2B).

**Figure 2:**
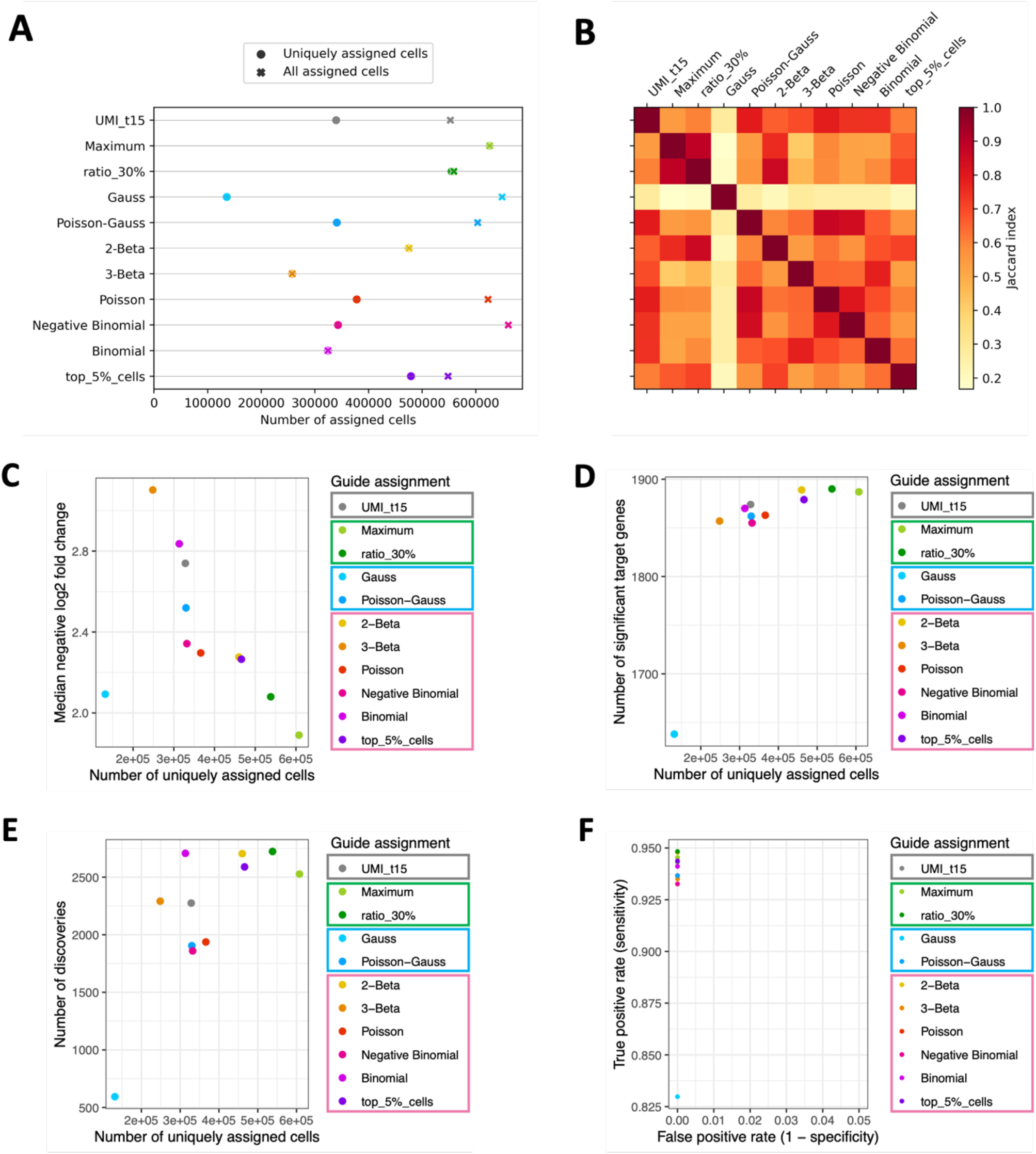
Results for the K562 cells of the Replogle screen. A: Number of total assigned cells (crosses) and cells with a single gRNA assigned (filled circle) per method. B: Assignment similarity heatmap showing the Jaccard index of the assignments per method pair. Only cells with one assigned gRNA (uniquely assigned cells) were considered. C-E: Median negative log2 fold change of the target gene per gRNA (C), number of significant targets (D) and number of total discoveries (E) (y-axis) vs. the number of uniquely assigned cells per method (x-axis). The number of total discoveries was calculated for a subset of 40 selected gRNAs and 5000 response genes (see Supplementary Information for details). F: True positive rate (y-axis) vs. false positive rate (x-axis) of all methods. Coloured boxes in the legend mark the four guide assignment groups as shown in Fig. 1.

For every assignment we then quantified differentially expressed genes in cells assigned to a targeting gRNA compared to control cells (assigned to non-targeting gRNAs) - also called discoveries in the following. Based on this, we evaluated the effectiveness in downregulating the target gene per gRNA as a positive control as well as the number of total and false discoveries for every assignment method (Supplementary Methods). In general, methods with a smaller number of uniquely assigned cells tended to have a stronger downregulation of the targeted gene per gRNA (higher negative log2 fold change, Fig. 2C), indicating a more restrictive assignment of only high-confidence cells. In contrast, methods with larger numbers of uniquely assigned cells identified more target genes with significantly changed expression, likely due to the increased sample size for statistical testing (Fig. 2D).

While most methods in crispat learn an optimal threshold from the data, a subset requires a user-defined threshold as input (Supp. Table 1). For these methods we used crispat to test a wide range of values and compared how the threshold choice affects the performance metrics (Supp. Fig. 2). The choice of the threshold strongly influenced the performance of the method, with thresholds resulting in a larger number of uniquely assigned cells leading to a lower median target downregulation in the assigned cells. This resulted in a trade-off between sample size and effect size when testing for significant changes in gene expression.

When inspecting the number of discoveries beyond the target gene we observed a similar positive relationship between the number of uniquely assigned cells and the number of discoveries (Fig. 2E). Overall, the number of discoveries varied between 594 and 2,769 depending on the choice of the guide assignment method. While mostly driven by the difference in the number of uniquely assigned cells, we also noted differences of more than 800 discoveries between methods with similar cell counts.

Another important criterion in choosing a suitable guide assignment is the rate of false discoveries associated with a given assignment method. To quantify this, we tested for differential expression in cells assigned to a given non-targeting gRNA against the remaining control cells assigned to all other non-targeting gRNAs. Here, we would expect no differences in the gene expression and the number of discoveries in this experiment can thus provide an indication of the false positive rate. All methods identified at most two false discoveries for non-targeting control gRNAs (out of 86,880-104,760 tested associations) resulting in a small false positive rate (number of false discoveries out of all tested gene - non-targeting gRNA pairs, Fig. 2F), and the differential expression test seemed to be similarly well calibrated for all assignments on this data set (Supp. Fig. 4).

A notable outlier in the method comparison was the assignment based on a Gaussian mixture model, which despite very few cells with a unique guide assignment showed a very low target downregulation in the assigned cells (Fig. 2C). As this method is routinely applied when using 10X Genomics Cell Ranger for data processing we investigated what is causing the bad performance. We noted that the thresholds inferred by this model were very low for all gRNAs, leading to more than 600,000 cells with at least one assigned gRNA and a majority of cells having more than one gRNA assigned (Fig. 2A). This is in contrast to the expectations based on the experimental MOI of the screen and typical downstream analyses focus on uniquely assigned cells. To investigate to what extent these assignments of a single cell to many gRNAs could still be used we repeated the differential expression analysis including also cells with multiple gRNAs assigned. In this analysis, the Gaussian mixture model resulted in the highest number of discoveries of all assignment methods (Supp. Fig. 5A) but with an excessive number of false discoveries (n=3,123) (Supp. Fig. 5B). These results indicate that the assignment by the Gaussian mixture model is too lenient, with very low thresholds not being able to discern background contamination. We hypothesized that this problem is especially severe for large-scale CRISPR screens with a high number of different gRNAs, where many cells have zero counts for a given gRNA and are included in the mixture model. To verify this we re-evaluated the method on semi-synthetic guide counts reducing the number of gRNAs in the Replogle data by aggregating different gRNAs into a smaller set of 500 or 86 gRNA groups, resulting in a performance of the Gaussian mixture model that was comparable to other methods (Supp. Fig. 5C,D). This suggests that the method requires adaptations for CRISPR screens with many gRNAs. We observed that a Gaussian mixture model fitted only on non-zero gRNA count values can remedy this problem to some extent, resulting in a strong increase in the number of significant target genes and total discoveries and a more similar performance to the other mixture models (Supp. Fig. 5E,F,G). This is therefore included as an option in crispat.

Another consideration for the choice of a guide assignment strategy is its run time, both for the assignment as well as downstream analyses. For the guide assignment, the Poisson mixture model as well as an adaptation thereof using a negative binomial distribution had the longest run time of up to 170 hours without parallelization (Supp. Fig. 6A). It is important to note here however that the implementation in crispat allows parallelization across gRNAs with a run time of around 4.5 minutes per gRNA. When using parallelization with 32 processes for the three slowest methods, each assignment method can be run within 12 hours for the K562 cells of the Replogle screen (Supp. Fig. 6A). The run time for the downstream analyses using the SCEPTRE package^18^ was mostly driven by the number of assigned cells (Supp. Fig. 6B-D) such that methods achieving the same power using fewer cells would be advantageous in terms of both compute time and memory requirements.

Finally, we investigated to what extent a suitable guide assignment strategy has to be chosen in a dataset-specific manner. For this we first used crispat on a second cell line (RPE1 cells) from the same study^5^ and compared the relative performance of the guide assignment methods for the two cell lines following the same analysis steps as outlined above for the K562 cells. We again tested a range of different thresholds for the assignment methods that require a user-defined threshold and selected the optimal threshold with respect to the number of significant target genes (Supp. Fig. 7). Similar to the results for the K562 cells, we observed that methods with a higher number of uniquely assigned cells had a less effective target gene downregulation but identified more target genes as well as other genes (total discoveries) as significantly differentially expressed (Supp. Fig. 8). Overall, our analyses revealed a very high consistency between the two cell types in the relative performance of individual assignment methods across metrics regarding the assignment quality, i.e. target down-regulation, number of significant differentially expressed target genes and number of discoveries (Supp. Fig. 9A), as well as regarding the efficiency, i.e. number of uniquely assigned cells and run time (Supp. Fig. 9B).

We next asked whether different technologies and dimensions of the CRISPR screens influence the suitability and performance of individual methods. To test this, we used crispat on a CRISPRi screen with only 86 gRNAs and compared two different read-out strategies (whole transcriptomics and TAP-seq)^4^. Again, suitable thresholds have been selected for the methods that require a user-defined threshold based on the number of significant target genes in the two data sets (Supp. Fig. 10, 11). In general, differences across guide assignment methods were less pronounced in these screens with similar numbers of assigned cells per method as well as higher Jaccard indices across methods (Supp. Fig. 12A,B and 13A,B). As on the data by Replogle et al., we observed that methods with more uniquely assigned cells showed a less pronounced target gene downregulation in the assigned cells (Supp. Fig. 12C and 13C). Differences in the number of significant target genes and total discoveries were less pronounced between methods than in the Replogle et al. data (Supp. Fig. 12D-F and 13D-F). In general, the number of uniquely assigned cells of a method showed a positive relationship to the total number of discoveries as in the data by Replogle et al., with exceptions for methods assigning too many cells with low target downregulation, as e.g. the Maximum approach on the TAP-seq data (Supp. Figure 13). Overall, methods performing well in the genome-scale screen were not necessarily a good choice for this data, highlighting the need for data-specific comparison of different guide assignment methods.

## Discussion

Guide assignment is a crucial step in single-cell CRISPR screens since downstream analyses rely on the assigned gRNA labels. As shown in this study, the number of assigned cells as well as the number of total discoveries can vary by a factor of 4 depending on the chosen assignment method. Therefore, we created the python package crispat, which facilitates the comparison of 11 different assignment methods, and enables users to choose a suitable method based on different performance metrics, including the true and false positive rate of each method. While the true positive rate can be estimated based on known target genes for CRISPR interference screens, other CRISPR screens might require different criteria for the choice of the assignment method. For example, in screens with unknown target genes, e.g. enhancer screens, or no changes in target gene expression, e.g. CRISPR knockout screens, the true positive rate cannot be directly calculated. Here, an alternative could be to consider the number of total discoveries instead, possibly calculated on a small subset of gRNA-gene pairs. While such evaluation metrics can be calculated reasonably fast for small data sets (5-8 mins per assignment method using SCEPTRE on the TAP-seq data of Schraivogel et al. with 72 genes, 56 targeting gRNAs and around 22,000 cells), it requires substantially more time and memory on large genome-scale screens such as the one by Replogle et al. In these cases, we recommend the differential expression analysis on a subset of targeting gRNAs and select a good guide assignment method based on the true and false positive rate in this data subset.

While we here focus on the step of guide assignment, it is important to note that choices in the analysis strategy before and after guide assignment will have a strong impact on the overall results that can be obtained with crispat. Since all assignment methods in crispat are based on the gRNA-cell count matrix, choosing a suitable alignment strategy is crucial. For example, it has been shown previously that mutations can arise in gRNAs which can impair downstream analyses^19^. Alignment strategies that account for such mutations could serve as alternative input to crispat. Following guide assignment with crispat, the learnt assignments can serve as input to a wide range of different downstream analysis, including differential expression analysis with SCEPTRE, Seurat or scanpy^16–18^ or dimension reduction ^20,21^.

Finally, all assignment methods in crispat are based only on information from the gRNA counts. Interesting future work would be to additionally leverage other information about the successful perturbation in a cell such as markers for CRISPR efficiency or information from the mRNA information as proposed by GLM-EIV^13^ or MIMOSCA^22^. While such methods could yield more accurate assignments, future work is still needed to explore when guide assignment can benefit from including such information and to what extent statistical double dipping might be a problem. For this, crispat could be a starting point for a larger benchmarking study to derive general recommendations on the question of guide assignment in single cell CRISPR screens and disentangle the impact that various factors might have on the guide assignment, including capture technology and sequencing depth, number of target genes, coverage, multiplicity of infection and number of non-targeting versus targeting gRNAs.

## Acknowledgements

We thank Timothy Barry for valuable discussions about the SCEPTRE package, as well as Joseph Replogle for providing additional data from their analysis. This work was funded by the Deutsche Forschungsgemeinschaft (DFG, German Research Foundation) – 540147573. The authors acknowledge support by the state of Baden-Württemberg through bwHPC and the German Research Foundation (DFG) through grant INST 35/1597-1 FUGG, as well as the data storage service SDS@hd supported by the Ministry of Science, Research and the Arts Baden-Württemberg (MWK) and the German Research Foundation (DFG) through grant INST 35/1503-1 FUGG.

## Contributions

JB implemented the methods, analyzed the data and generated the figures

JB and BV wrote the manuscript

BV conceived and supervised the project

