## Supplementary Information for "Guide assignment in single-cell CRISPR screens using crispat"

### Supplementary Methods

#### Guide assignment strategies

In crispat, 11 different guide assignment methods have been implemented that can be grouped into 4 groups based on whether information is shared across gRNAs, cells or both. An overview of all our methods is shown in Table 1.

| Method | Group | Data type | Need user-specified threshold? | Batch | Total number of gRNAs per cell | MOI | References / New |
|---|---|---|---|---|---|---|---|
| UMI_t | Indepen-dent | counts | yes | not considered | not used | both | Schraivogel[1] Geiger-Schuller[2]* Adamson[3]** Frangieh[4]*** |
| Maximum | Across gRNAs | counts | no | not considered | not used | low | Papalexi[5]****, Jiang[6]***** |
| Ratio_X% | Across gRNAs | frequen-cies | yes | not considered | included in frequencies | low | Datlinger[7] |
| Gauss | Across cells | counts | no | applied per batch | not used | both | 10X Genomics Cell Ranger (version 7.2.0) |
| Poisson-Gauss | Across cells | counts | no | not considered | not used | both | Replogle[9] |
| 2-Beta | Across gRNAs & cells | frequen-cies | no | applied per batch | included in frequencies | low | new |
| 3-Beta | Across gRNAs & cells | frequen-cies | no | applied per batch | included in frequencies | low | new |
| Poisson | Across gRNAs & cells | counts | no | batch as a covariate | included as a covariate | both | Barry[10] |
| Negative Binomial | Across gRNAs & cells | counts | no | batch as a covariate | included as a covariate | both | new |
| Binomial | Across gRNAs & cells | counts | no | batch as a covariate | distribution parameter | low | new |
| top_X%_ cells | Across gRNAs & cells | frequen-cies | yes | not considered | included in frequencies | low | new |

**Table 1: Overview over all guide assignment methods.**
* in combination with a threshold on the ratios
** in combination with a threshold on the reads
*** relaxed for guides with a high number of reads
**** additional constraints to remove ambiguous cells
***** additional filtering of ambiguous cells with counts from many gRNAs and post-hoc refinement using negative cells with known absence of a gRNA

In the following we give a more detailed description of the individual methods in each group:

*Independent*

- **UMI threshold (UMI_t)**: This method uses a threshold on the number of UMI counts, assigning cells to all gRNAs that have at least as many UMI counts as specified by the threshold *t* in the cell. Due to its simplicity, this method has been widely used in the literature with various thresholds[1–4]. However, users have to define a suitable threshold for applying this method, which can depend on various factors such as the capture technology and sequencing depth of the screen.

*Across gRNAs*

- **Maximum**: This method assigns each cell the gRNA with highest UMI count in this cell. If there is no unique maximum in a cell, no gRNA is assigned. Since this approach assigns at most one gRNA to each cell, it is specifically designed for low MOI CRISPR screens.
- **Relative frequency threshold (Ratio_X%)**: This method assigns for each cell the gRNA with highest counts in this cell if its counts comprise at least *X%* of the total gRNA counts in this cell. As for the maximum approach, this method is designed for low MOI CRISPR screens and is for example the default assignment method for low MOI screens in the R package SCEPTRE[11]. Like the UMI threshold method, it relies on a user-defined threshold as input.

*Across cells*

- **Poisson-Gaussian mixture model (Poisson-Gauss)**: For every gRNA, this method fits a Poisson-Gaussian mixture model on the log2-transformed non-zero UMI counts of this gRNA over all cells across all batches. Next, all cells for which the probability of observing the guide counts from the Gaussian component is higher than for the Poisson (background) component are assigned to this gRNA. This approach was first proposed by Replogle et al.[9] Since no runnable code was provided in their paper, we re-implemented the model. While the method is applied across all batches, users can account for differences in sequencing depths by downsampling gRNA counts to a common value per cell. As this choice can be very dataset specific, crispat by default uses no downsampling, resulting in some but overall small differences to the original assignments reported by Replogle et al.[9](Supp. Fig. 13A).
- **Gaussian-Gaussian mixture model (Gauss)**: For every gRNA, this method fits a Gaussian-Gaussian mixture model on the log10-transformed UMI counts of this gRNA with a pseudocount of 1 over all cells in a batch. In crispat we implemented a Gaussian mixture model based on 10X Genomics Cell Ranger (version 7.2.0) assignment approach in which either variational inference via pyro or an EM algorithm based on sklearn (as in 10X Genomics Cell Ranger) can be selected. For

all analyses shown in this paper, the EM inference method has been selected. Additionally, we provide the option to fit the model only on non-zero counts which we found to be beneficial for large-scale screens (Supp Fig. 4F).

*Across cells and gRNAs*

- **2-Beta mixture model (2-Beta)**: Like the Ratio_X% approach, this method calculates for every cell the relative frequency of a gRNA as the ratio of its counts over the total number of gRNA counts in a cell and assigns a cell to the most abundant gRNA in this cell if its relative frequency is a above a certain threshold. In the Ratio_X% method, this threshold has to be chosen manually. By modeling the relative frequencies of the most abundant gRNA across all cells as a mixture distribution, the Beta mixture models can directly learn a suitable threshold. In the 2-Beta mixture model, this aims at distinguishing cells with a high relative frequency of the most abundant gRNA (higher mode of the mixture distribution, most guide counts in a cell coming from a single gRNA, providing evidence for a transfection of this cell with the gRNA) from cells with a low ratio of the most abundant gRNA (lower mode of the mixture model, no single gRNA stands out clearly, counts of the most abundant gRNA are in a similar range to counts from all other gRNAs). The model is fitted across all cells from a given batch to determine a threshold on the relative frequency based on where the two Beta distributions intersect. This results in one threshold per batch (without distinguishing between gRNAs), which is then used as X in the Ratio_X% approach. Since this method is based on the Ratio_X% approach and looks for single gRNAs in a cell that stands out compared to the other gRNAs in that cell, it is designed for low MOI CRISPR screens.
- **3-Beta mixture model (3-Beta)**: This method uses the same approach as the 2-Beta model but using a 3-component Beta mixture and defining the threshold X as the intersection of the two highest components. The third component can thus capture cells infected with two gRNAs.
- **Latent variable Poisson generalized linear model** (**Poisson**): For every gRNA, this method fits a Poisson mixture on the UMI counts of this gRNA across all cells with mean $\lambda = e^{\beta_0 + \beta_1 p_c + \beta_2 b_c + log(s_c)}$ with $\beta_0 \in R$, $\beta_1 \in R^+$, $\beta_2 \in R^n$, a latent perturbation state variable $p_c \in \{0, 1\}$ and cell-specific covariates (total number of gRNA counts per cell $N_c$ and one-hot encoded batch $b_c$ for $n$ batches). This approach was proposed by Barry et al.[10] and is implemented in the R package SCEPTRE[11]. For larger flexibility and due to memory limitations of SCEPTRE for the application on the data by Replogle et al.[9], we implemented a custom version of this mixture model as part of crispat, which yields very similar assignments (Supp. Fig. 13B). Newer versions of SCEPTRE based on Nextflow will accommodate a direct application of their algorithm.
- **Latent variable Negative Binomial generalized linear model (Negative Binomial)**: This method is a modified version of the Poisson method using a Negative Binomial distribution instead of Poisson distribution. The overdispersion is learnt as an additional parameter in the model.
- **Latent variable Binomial generalized linear model (Binomial)**: For every gRNA, this method fits a binomial distribution $B(N_c, \theta_c)$ with $N_c$ being the total number of

gRNA counts per cell and $\theta_c = sigmoid(e^{\beta_0+\beta_1 p_c+\beta_2 b_c})$ with $\beta_0 \in R$, $\beta_1 \in R^+$, $\beta_2 \in R^n$, latent perturbation state $p_c \in \{0, 1\}$ and one-hot encoded batch $b_c$.

- **Quantile approach (top_X% cells)**: For every gRNA, this method chooses the top X% of cells with the highest gRNA counts, excluding cells with zero counts for the gRNA. This is related to the Ratio_X% method with a gRNA-specific threshold that depends on the relative frequencies of this gRNA in other cells. This method requires a user-defined threshold as input which determines what percentage of cells will be assigned for every gRNA.

**Data set from Replogle et al.**

For our benchmark study, we used data from a single-cell CRISPR interference screen by Replogle et al.[9] in which 2,285 common essential genes in K562 and RPE1 cells were perturbed. In the original study, the authors have used 10X Genomics Cell Ranger to align the sequencing reads to the reference genome and to the gRNA library. We downloaded the output from the filtered_feature_bc_matrix directory which includes 3 files for each batch (barcodes.tsv.gz, features.tsv.gz and matrix.mtx.gz) from the following link: https://plus.figshare.com/articles/dataset/_Mapping_information-rich_genotype-phenotype_landscapes_with_genome-scale_Perturb-seq_Replogle_et_al_2022_MTX_files/20127869/1

We used scanpy (version 1.9.3) to read in the data per batch and created an AnnData object combining all batches (AnnData version 0.9.2), containing 686,464 cells, 36,601 measured transcripts and 2,283 gRNA pairs across 48 batches for the K562 cells and 616,184 cells, 36,601 measured transcripts and 2,679 gRNA pairs across 56 batches for the RPE1 cells. The screen by Replogle et al. was designed such that each lentivirus contains two gRNAs for the same target gene (multiplexed CRISPRi library). We summed the gRNA counts of the two gRNAs from a pair and applied the gRNA assignment on the gRNA pair level. Since several guide assignments are applied on the relative counts per cell, this step is necessary to distinguish cells with single infections from double infected cells in these methods.

**Data set from Schraivogel et al.**

As a comparison, we applied the guide assignment methods to single-cell CRISPR interference screens in K562 cells from Schraivogel et al.[1] in which 14 regions (promoters and enhancers of known target genes) were targeted with 4 gRNAs each (56 targeting + 30 non-targeting control gRNAs). Gene expression and gRNA count matrices for whole transcriptome Perturb-seq and targeted Perturb-seq (TAP-seq) were downloaded from http://steinmetzlab.emgl.de/TAPdata/TAP.nods.RDS and http://steinmetzlab.embl.de/TAPdata/Whole.nods.RDS. The count matrices were read in with R and the gRNA counts were exported as a csv file to serve as an input for crispat. The resulting data contained 37,971 cells, 17,107 measured genes and 85 gRNAs for the whole transcriptomics screen and 21,977 cells, 72 measured genes and 86 gRNAs for the TAPseq screen. Since we observed many false discoveries for one specific non-targeting gRNA (non-targeting gRNA 22) across all assignment methods, we excluded this gRNA and its assigned cells for all downstream analyses.

**Evaluation of guide assignment methods**

Using crispat we applied all guide assignment methods to the gRNA count matrices for each of the four data sets and filtered the assignments to contain only cells with exactly one gRNA

(gRNA pair in the Replogle screen) assigned. To compare the concordance of assignments across methods, we calculated for every pair of methods the ratio of the number of cell-gRNA pairs found in both methods divided by the total number of cell-gRNA pairs found in a single method.

Next, we used the R package SCEPTRE (version 0.10.0)[11] to analyze how strongly the differences between guide assignments affect downstream analyses such as differential expression testing. In particular, we used the calibration check, power check and discovery analysis in SCEPTRE for every guide assignment obtained in crispat. With these steps, we test for false discoveries using negative control-target pairs, differential expression of known target genes and differential expression across all genes, respectively (see https://timothy-barry.github.io/sceptre-book/sceptre.html for details on the individual steps). Before creating the SCEPTRE objects for the two Replogle screens, we removed cells with less than 3000 total gene UMI counts and more than 20% or 11% mitochondrial genes in K562 and RPE1 cells, respectively, choosing the same cut-offs as in the original study by Replogle et al[9]. Otherwise, the following analysis and quality control parameters were used (defaults for variables not specified here):

- side = “both” (testing for up- and downregulation of gene expression)
- grna_integration_strategy = “union” for Replogle data set and grna_integration_strategy = “singleton” for Schraivogel data sets
- control_group = “nt_cells” (default in low MOI setting)
- multiple_testing_alpha = 0.05
- formula = log(response_n_nonzero) + log(response_n_umis) + log(grna_n_nonzero + 1) + log(grna_n_umis + 1) + response_p_mito + Batch + S_score + G2M_score) with number of non-zero genes (response_n_nonzero) and gRNAs (grna_n_nonzero), number of total gene UMIs (response_n_umis) and gRNA UMIs (grna_n_umis), percentage of mitochondrial gene counts (response_p_mito), batch and cell cycle scores (S_score, G2M_score) of each cell. For the Schraivogel data only one batch was present and thus the batch covariate excluded from the model, for the TAPseq data no cell cycle scores were included (due to the small number of measured genes, cell cycle scores cannot be calculated).
- n_umis_range = c(0,1) and n_nonzero_range = c(0,1) for the Replogle data sets since low quality cells were already removed before creating the SCEPTRE object with the thresholds described above. For the Schraivogel data set, default SCEPTRE parameters for the number of total UMIs and number of non-zeros were kept.
- n_nonzero_trt_thresh = 0 (such that perturbations in which the expression of a gene is zero across all cells in the perturbed condition are not excluded for testing)

Instead of running the guide assignment step of the SCEPTRE package, we used the results obtained from crispat as input and then followed the typical SCEPTRE pipeline. Due to high time requirements of the calibration and discovery analysis, for the Replogle data these steps were performed for a subset of 40 gRNAs, which were selected based on their number of assigned cells in the original study, choosing 10 gRNAs randomly from each 25% quantile. Additionally, for the calibration and discovery we selected 5000 random response genes in all data sets apart from the Schraivogel TAPseq screen which only contains the expression of 72 genes.

#### Experiments including cells with multiple assignments

For the K562 data set by Replogle et al.[9], we additionally evaluated the use of cells with more than one assigned gRNA. For this, SCEPTRE was used with the 'moi' parameter in the 'import_data' function set to 'high', testing for differences in gene expression compared to the control group given by all cells not assigned to a specific gRNA ('complement').

#### Experiments aggregating gRNAs into groups

To analyze the effects of the number of total gRNAs in the screen, we generated semi-synthetic data from the Replogle K562 screen by grouping guides into 500 or 86 groups of similar size and aggregated the gRNA counts per group. Guide assignment and discovery analysis were then performed for all assignment methods on these groups.

### Supplementary Figures

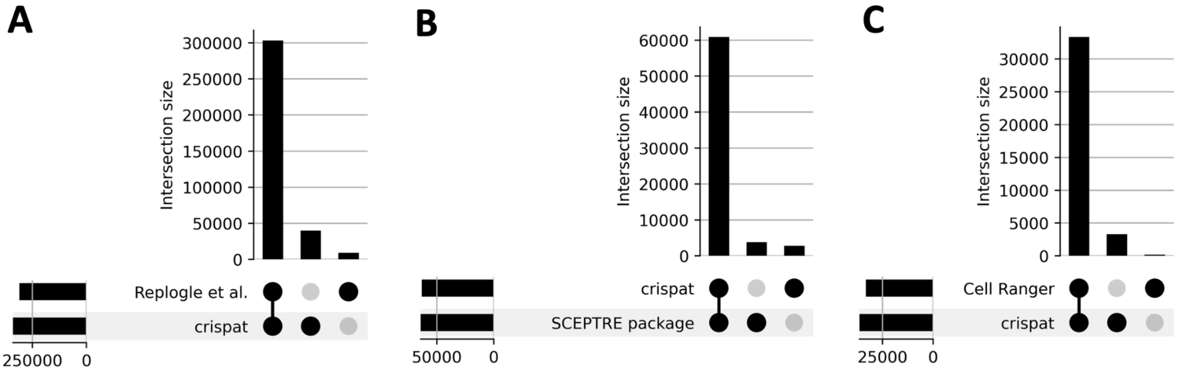


**Supplementary Figure 1: Assignment comparison in the K562 data set by Replogle et al. obtained from implementations in crispat to the original implementations.** A: UpSet plot comparing the crispat assignments using a Poisson-Gaussian mixture model as suggested by Replogle et al. to the original assignment labels reported in the study by Replogle et al. B: UpSet plot comparing the crispat assignments using a mixture model based on a latent variable Poisson generalised linear model (*Poisson*) as suggested by SCEPTRE vs. the assignments of the R package SCEPTRE (version 0.10.0). This comparison is shown for a data subset consisting of 40 batches and 50 gRNAs. C: UpSet plot for the crispat assignments using a Gaussian mixture model compared to the assignments obtained by Cell Ranger (version 4.0.0) on batch 1 (based on individual gRNAs thresholds by CellRanger provided by Replogle et al. instead of gRNA pairs). These three methods were re-implemented in crispat due to the lack of runnable code for (A), restrictions in the number of cells for (B) and an end-to-end pipeline without guide assignment as an individually executable step in (C).

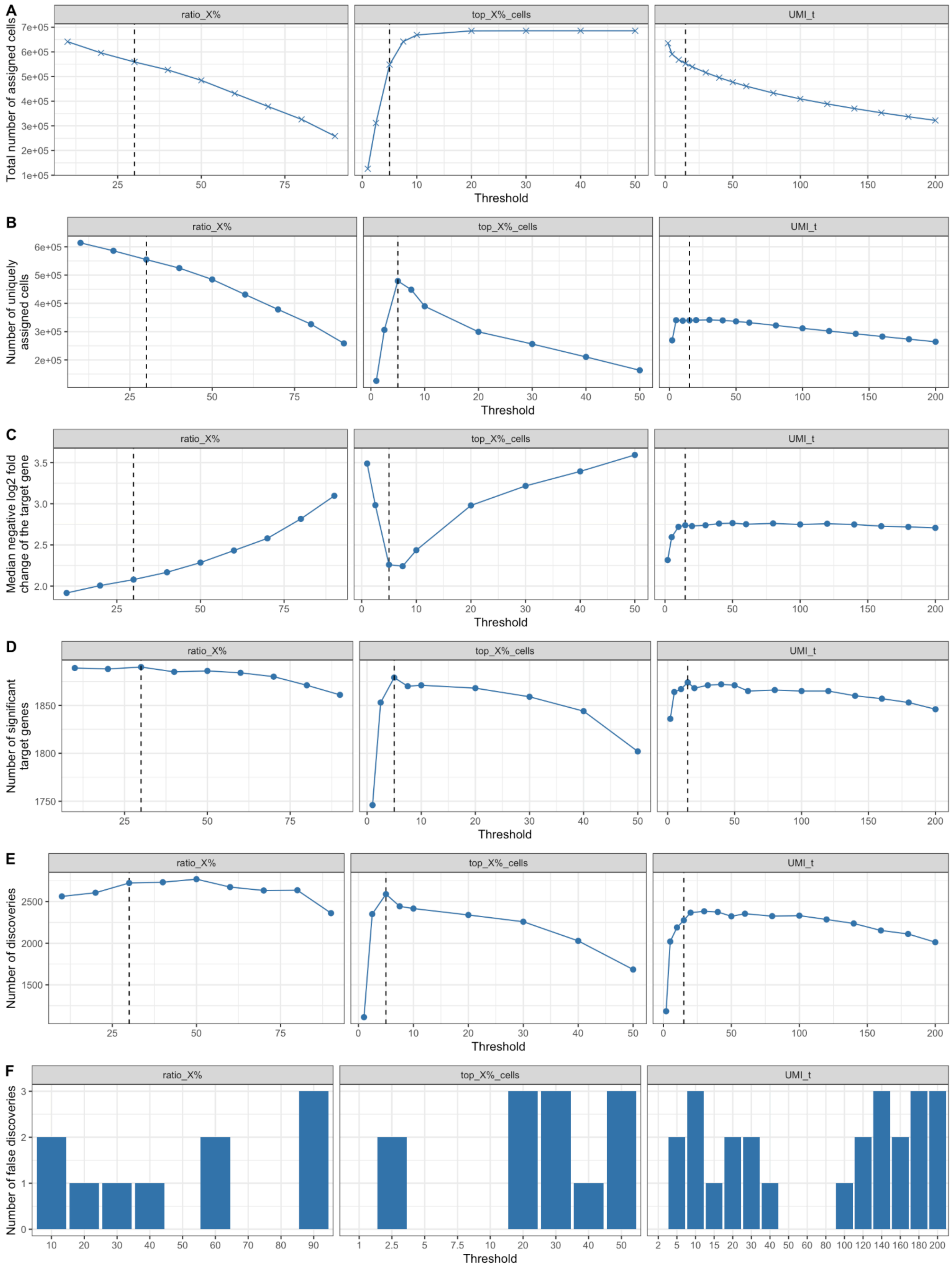


**Supplementary Figure 2: Effects of varying user-defined thresholds in the K562 data set by Replogle et al.** For each method where a threshold has to be defined (ratio_X% (left), top_X%_cells (middle) and UMI_t (right)), the total number of assignments (A), the number of cells with exactly one assigned gRNA (B), the median negative log2 fold change of the target gene (C), the number of significant target genes (D), the number of total discoveries (E) and the number of false discoveries (F) is shown over various thresholds (x-axis). The black dotted lines mark the chosen threshold per method based on the maximum number of significant target genes.

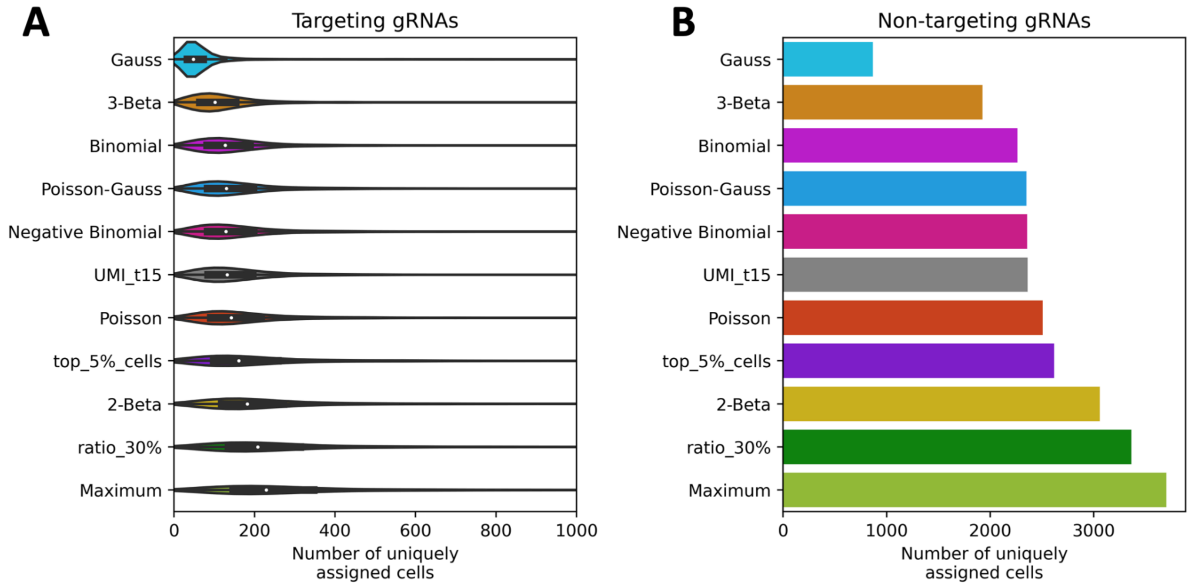


**Supplementary Figure 3: Number of uniquely assigned cells per gRNA in the K562 data set by Replogle et al.** Number of cells with exactly one gRNA assigned (x-axis) for every method (y-axis) distinguishing between cells assigned to targeting gRNAs (A) and non-targeting gRNAs (B).

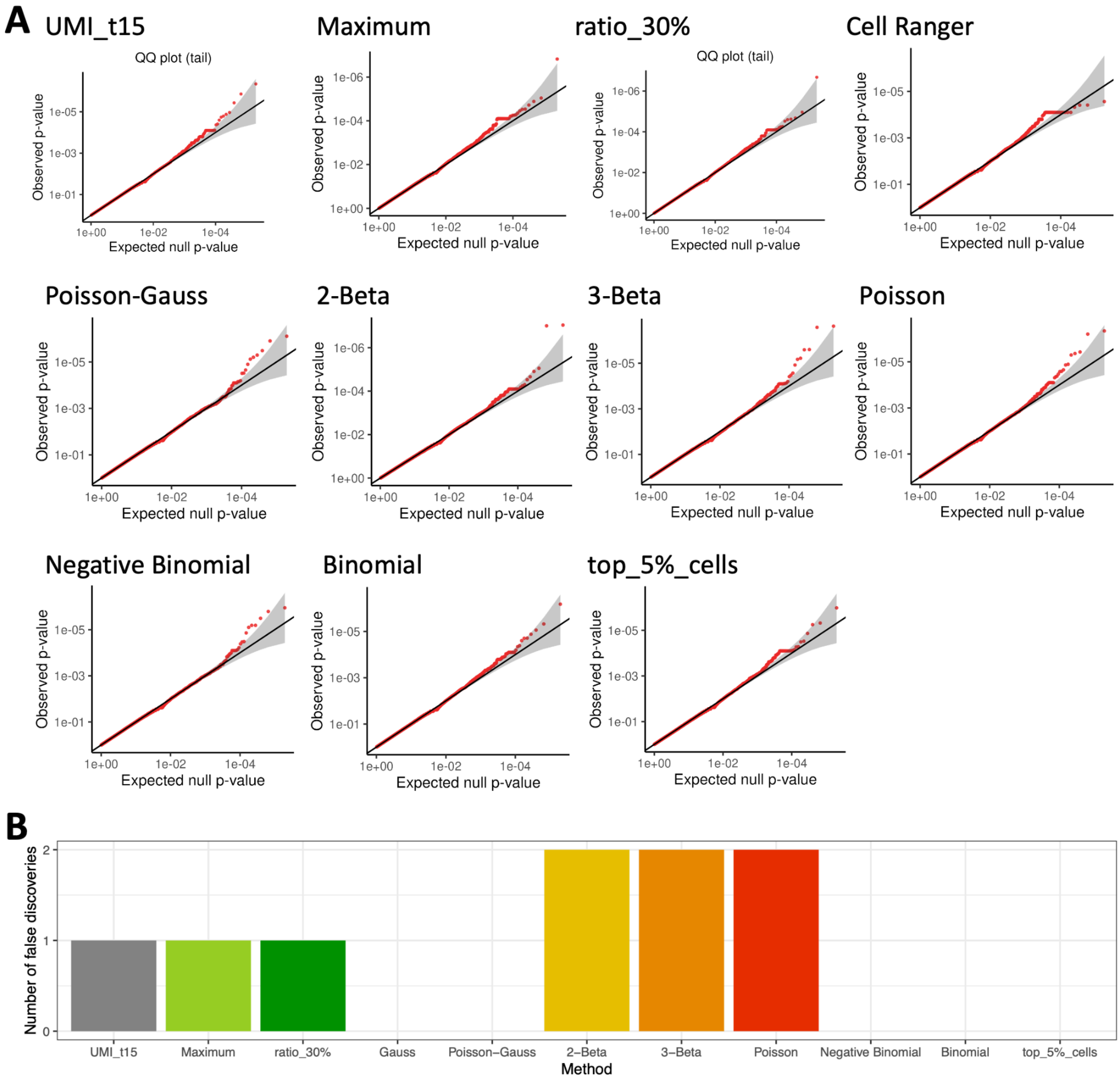


**Supplementary Figure 4: Calibration on the K562 data set by Replogle et al.** A: QQ-Plots for every assignment showing the p-values obtained in the calibration step of the SCEPTRE package in which the expression of cells assigned to one non-targeting gRNA is tested for differential expression compared to the other control cells. B: Number of false discoveries (y-axis) for every assignment method (x-axis) out of 86,880-104,760 tested associations.

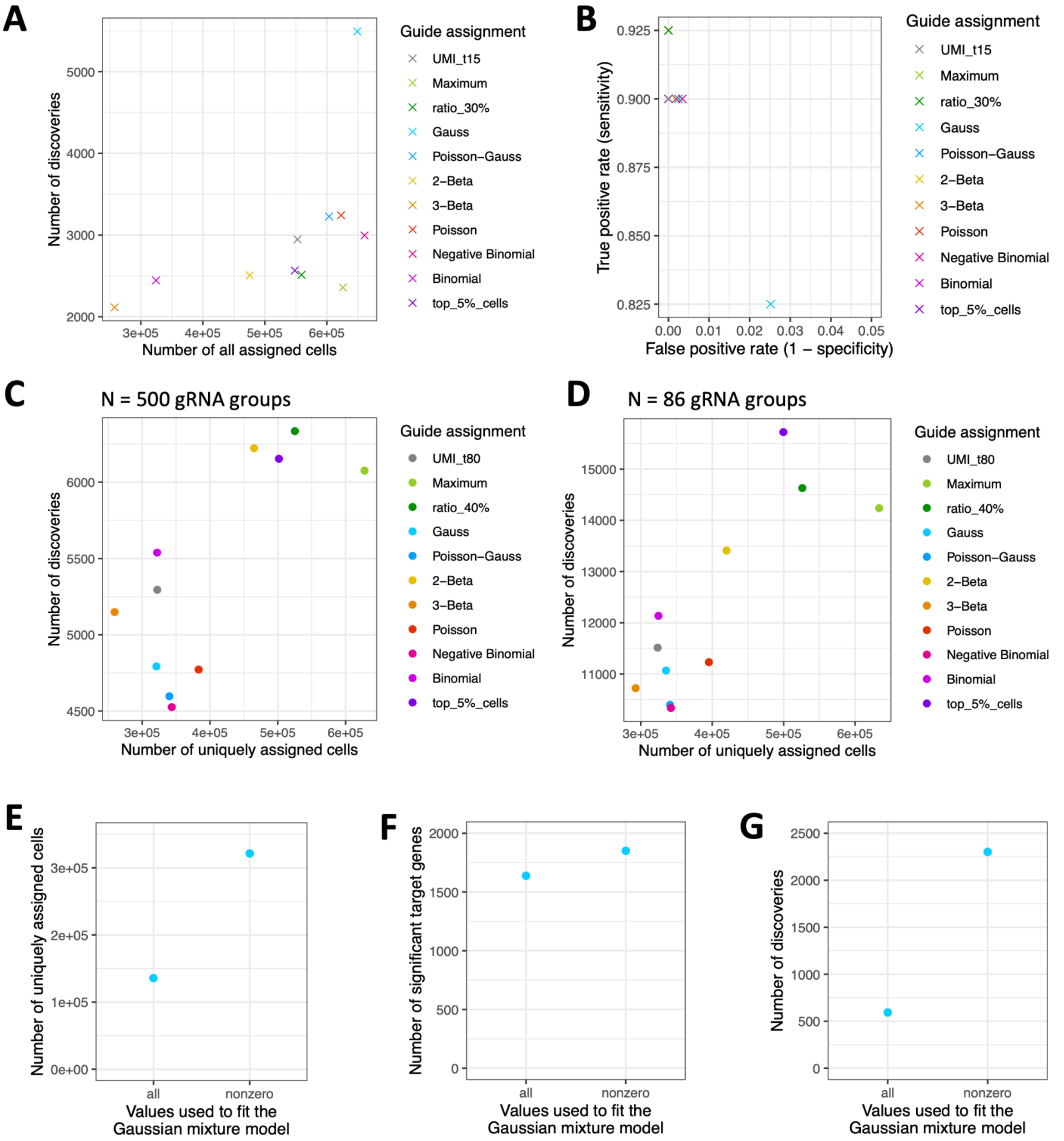


**Supplementary Figure 5: Investigation of the performance of the Gaussian mixture model in the K562 data set by Replogle et al.** A: Number of total discoveries for 40 selected gRNAs and 5000 genes (y-axis) vs. the number of assigned cells per method (x-axis) including cells with more than one gRNA assigned. C: False positive rate (x-axis) vs. true positive rate (y-axis) per method when cells with more than one gRNA are included as in A. C: Number of total discoveries for 40 selected gRNA groups and 5000 genes (y-axis) vs. the number of uniquely assigned cells (x-axis) when performing the guide assignment on 500 gRNA groups by combining the counts of random gRNAs with each other (see Supplementary Information for details). D: Same as in C for 86 gRNA groups (same number of gRNAs as in Schraivogel et al.). E/F/G: Comparison of fitting the Gaussian mixture model on all values vs. fitting on the non-zeroes regarding the number of cells with single gRNA assigned (E), number of significant target genes (F) and number of total discoveries (G). The number of false discoveries is not shown since they were zero in both cases. All analyses apart from panels A and B were conducted on uniquely assigned cells.

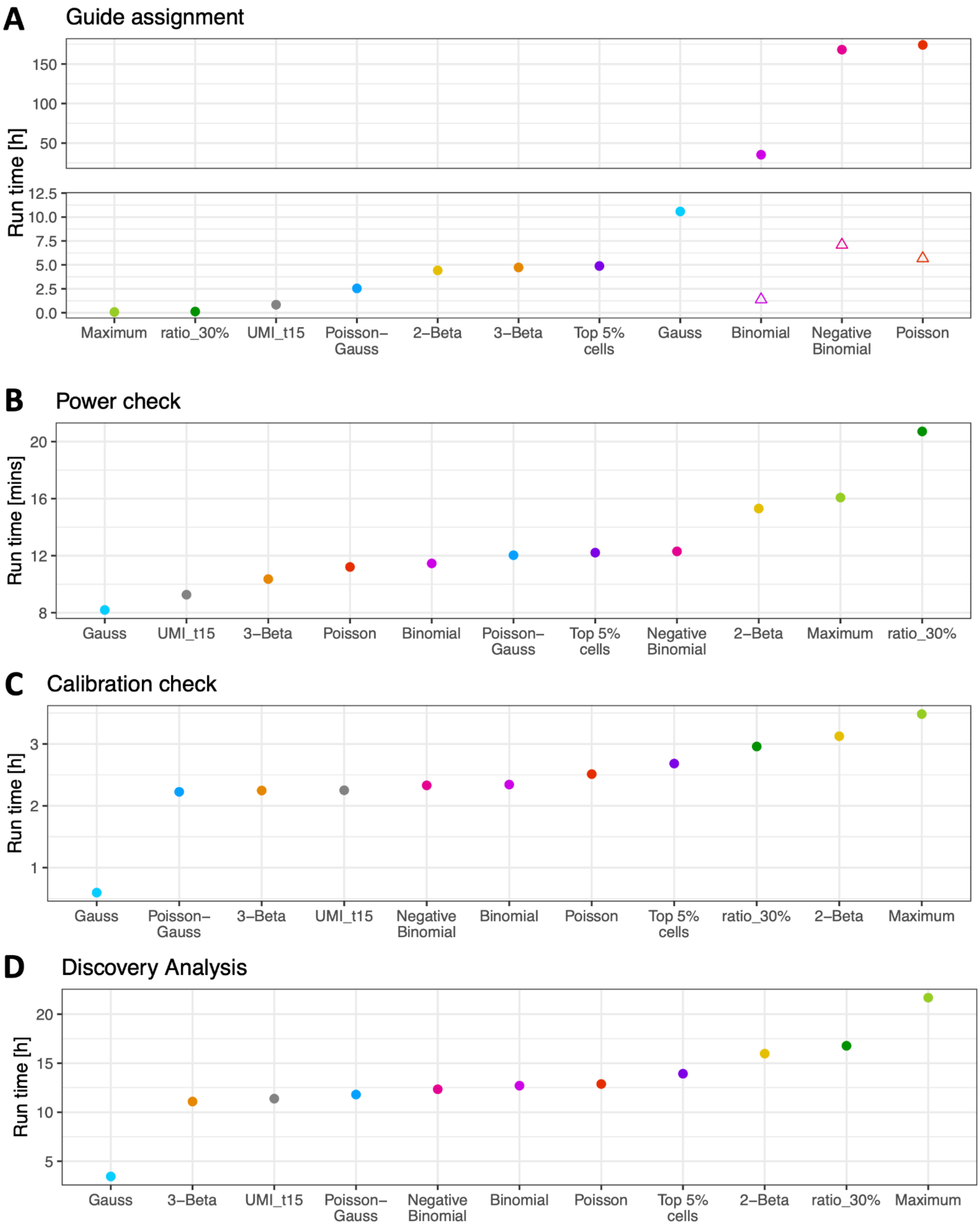


**Supplementary Figure 6: Run times for the analysis steps in the K562 data set by Replogle et al.** Run time (y-axis) of each guide assignment method (x-axis) for various steps of the analysis: guide assignment (A), power check (B), calibration (C), discovery analysis (D). Since the guide assignment takes the longest for the binomial, negative binomial and Poisson methods, crispat has an option to parallelize. Filled circles show unparallelized run times, triangles show the run time using 32 parallel processes. The calibration and discovery step of the SCEPTRE pipeline were run on a data subset consisting of 40 gRNAs, 5000 response genes and all cells assigned to a non-targeting gRNAs or one of these 40 gRNAs (see Supplementary Information for details).

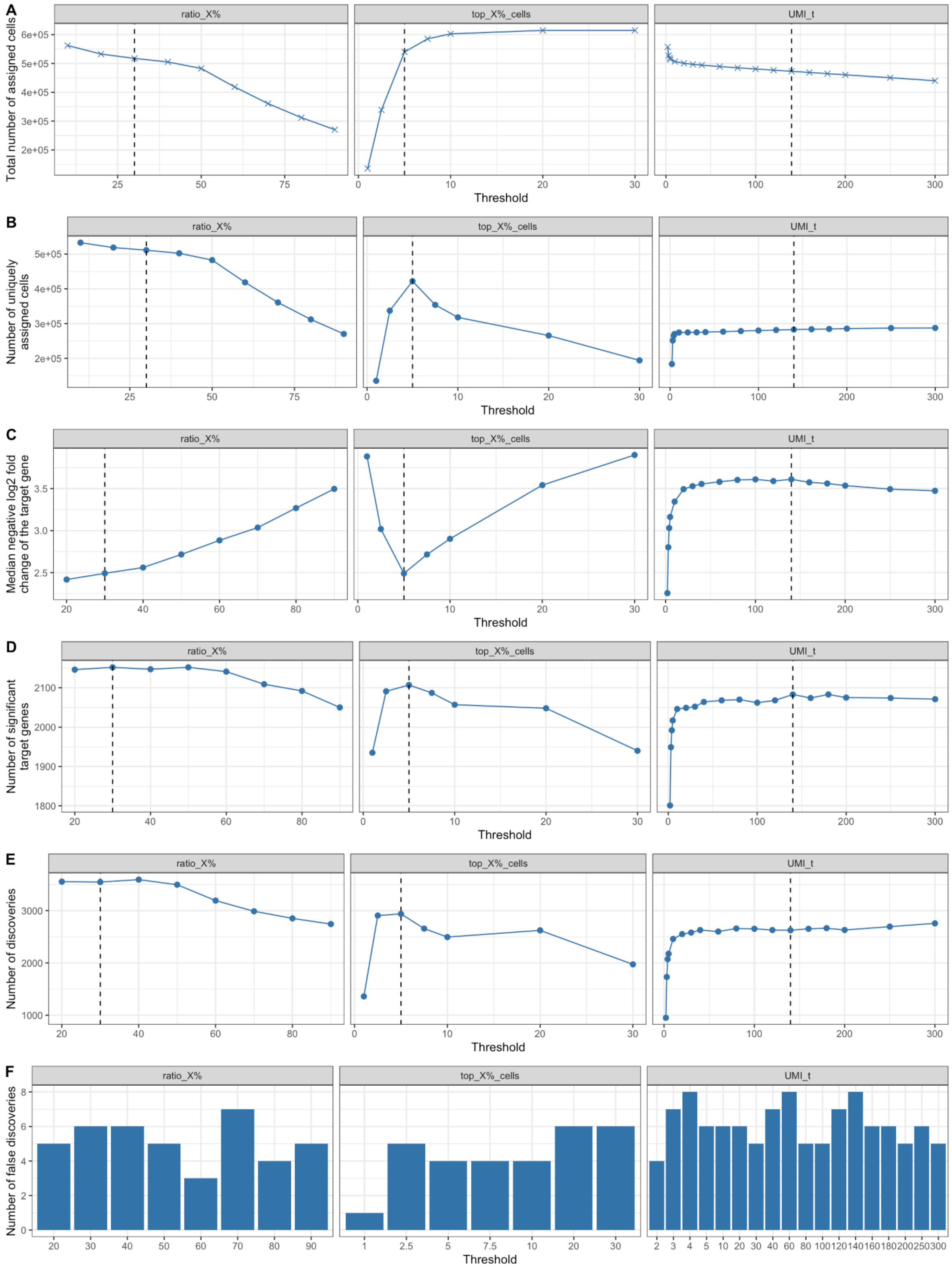


**Supplementary Figure 7: Effects of varying the threshold in the RPE1 data set by Replogle et al.** For each method where a threshold has to be defined (ratio_X% (left), top_X%_cells (middle) and UMI_t (right)), the total number of assignments (A), the number of cells with exactly one assigned gRNA (B), the median negative log2 fold change of the target gene (C), the number of significant target genes (D), the number of total discoveries on a data subset (E) and the number of false discoveries (F) is shown over various thresholds (x-axis). The black dotted lines mark the chosen threshold per method based on the maximum number of significant target genes.

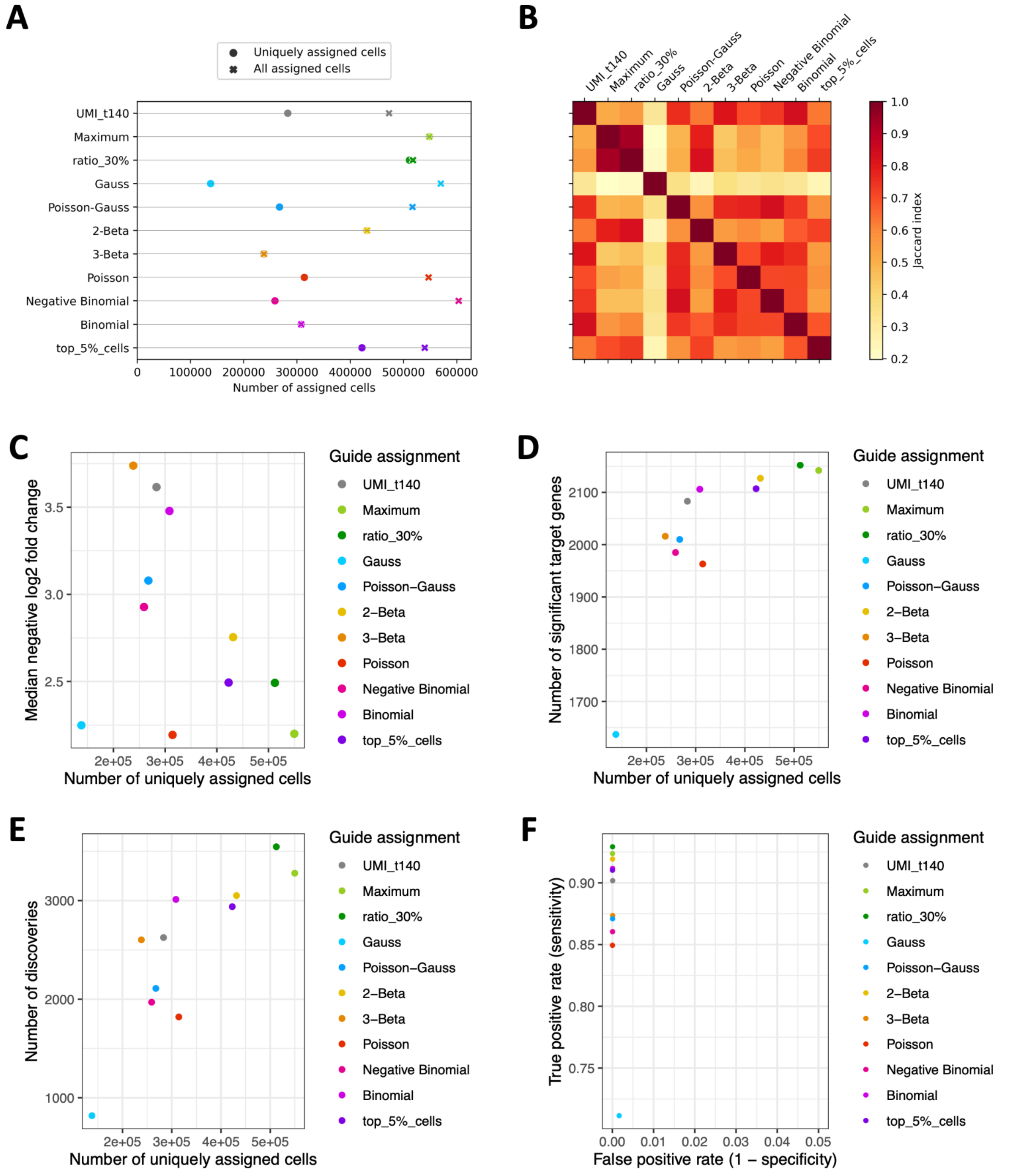


**Supplementary Figure 8: Method comparison on the RPE1 data set by Replogle et al.** A: Number of total assigned cells (crosses) and cells with a single gRNA assigned (filled circle) per method. B: Assignment similarity heatmap showing the Jaccard index of the assignments per method pair. Only cells with one assigned gRNA were considered. C-E: Median negative log2 fold change of the target gene per gRNA (C), number of significant targets (D) and number of total discoveries (E) (y-axis) vs. the number of uniquely assigned cells per method (x-axis). The number of total discoveries was calculated for a subset of 40 selected gRNAs and 5000 response genes (see Supplementary Information for details). F: True positive rate (y-axis) vs. false positive rate (x-axis) of all methods.

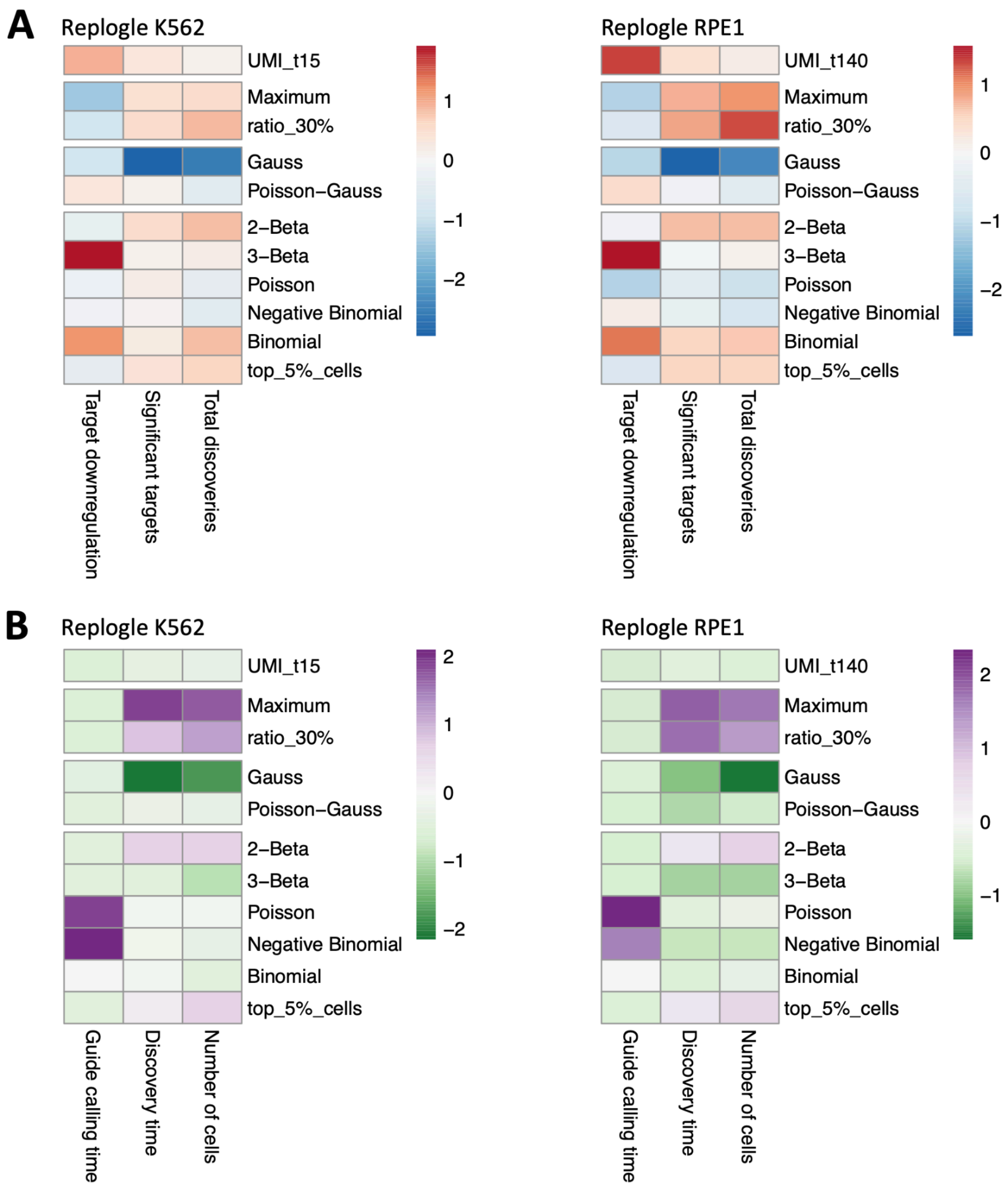


**Supplementary Figure 9: Comparison of guide assignment performance in K562 (left) vs. RPE1 (right) data sets by Replogle et al..** A: Comparison of three assignment quality metrics (x-axis) across guide assignment methods (y-axis): (i) negative median log2 fold change of the target gene, (ii) number of significant target genes, and (iii) total number of discoveries for 40 selected gRNAs on 5000 genes. The values of all three features have been z-normalised across methods. B: Comparison of three efficiency metrics (x-axis) across guide assignment methods (y-axis): (i) Number of assigned cells, (ii) run time for the (unparallelized) guide assignment, and (iii) run time for the discovery analysis using the SCEPTRE package.

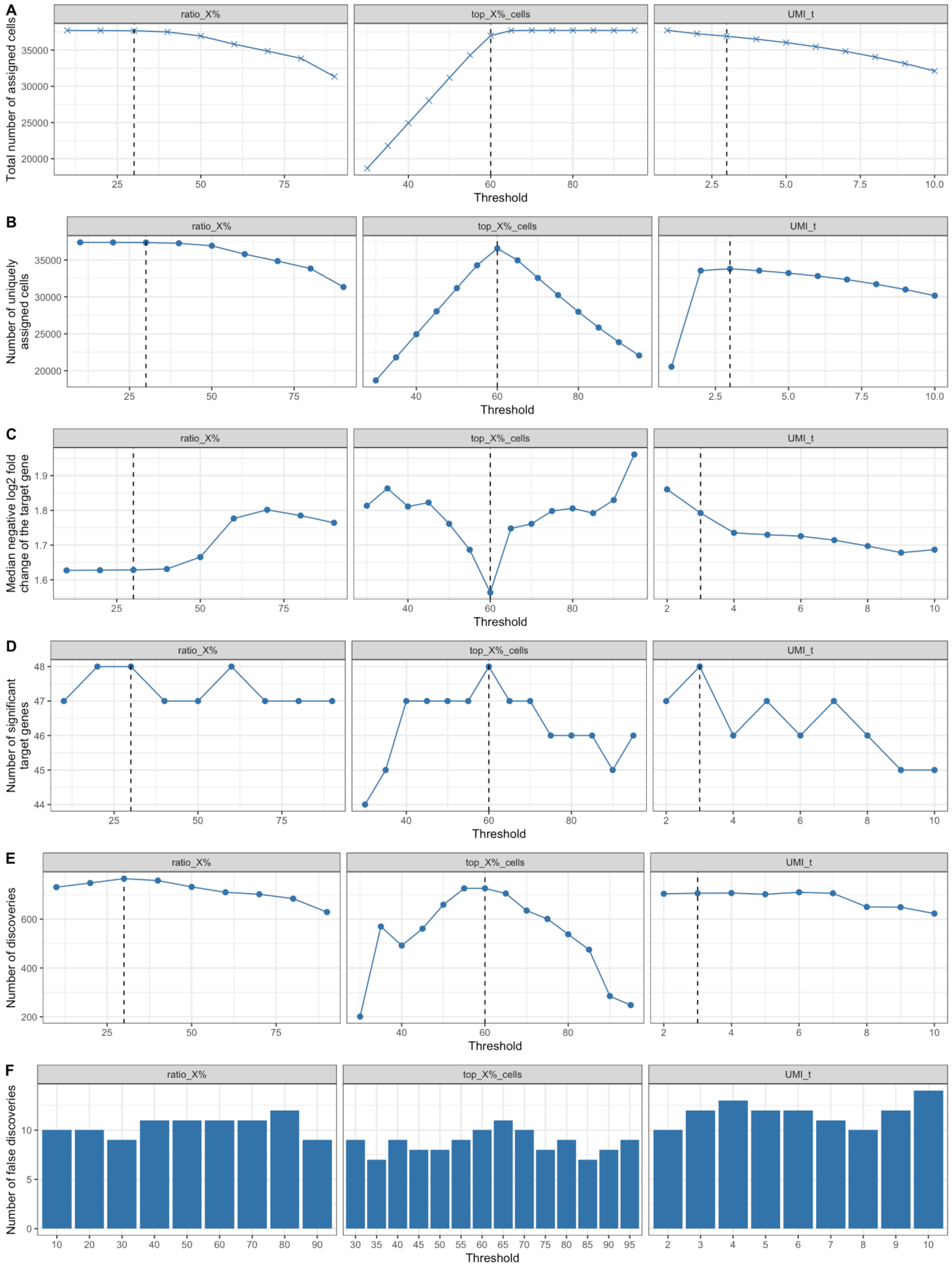


**Supplementary Figure 10: Effects of varying the threshold in the whole transcriptomics screen by Schraivogel et al.** For each method where a threshold has to be defined (ratio_X% (left), top_X%_cells (middle) and UMI_t (right)), the total number of assignments (A), the number of cells with exactly one assigned gRNA (B), the median negative log2 fold change of the target gene (C), the number of significant target genes (D), the number of total discoveries on a data subset (E) and the number of false discoveries (F) is shown over various thresholds (x-axis). The black dotted lines mark the chosen threshold per method based on the maximum number of significant target genes.

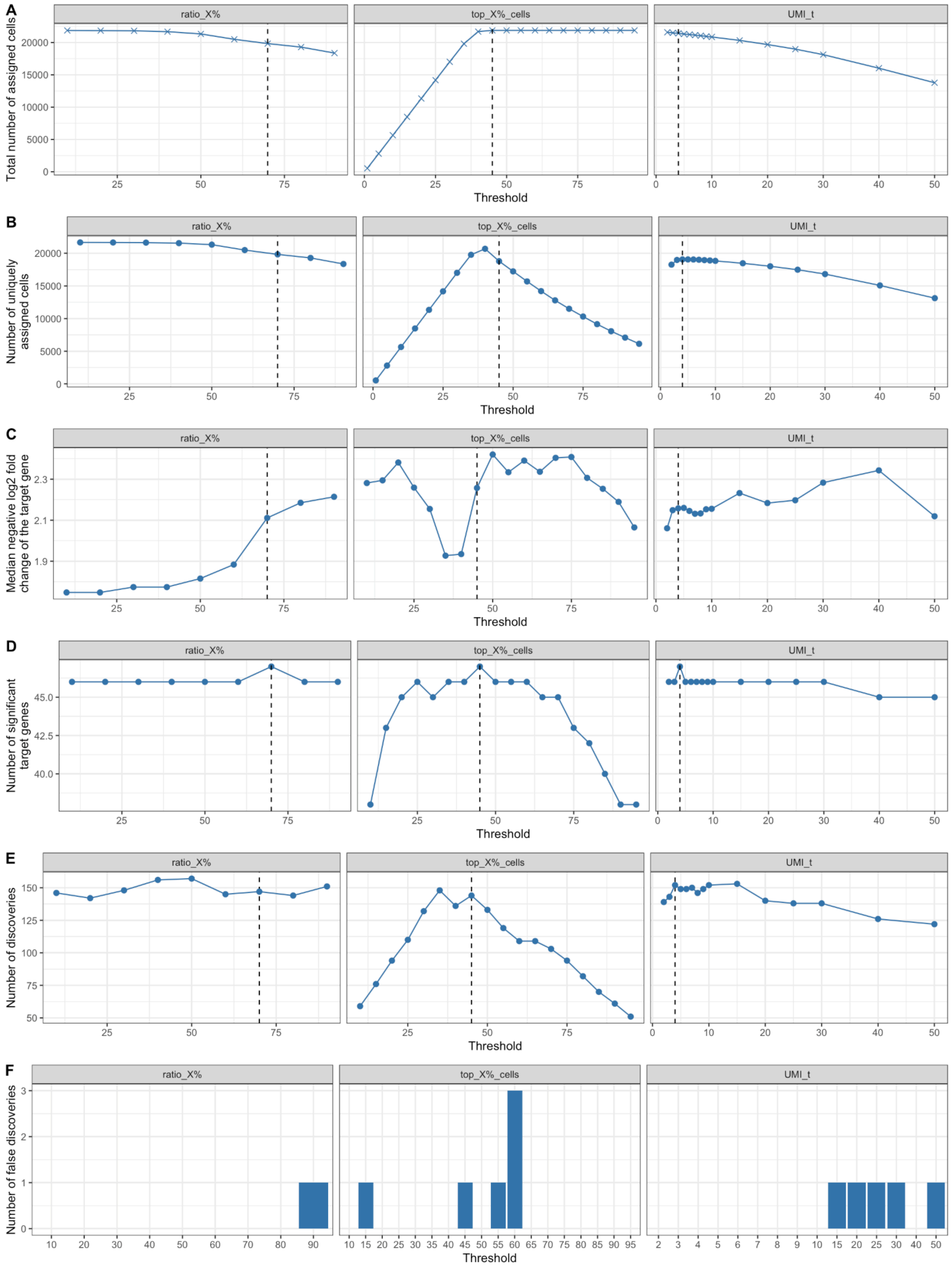


**Supplementary Figure 11: Effects of varying the threshold in the TAPseq screen by Schraivogel et al.** For each method where a threshold has to be defined (ratio_X% (left), top_X%_cells (middle) and UMI_t (right)), the total number of assignments (A), the number of cells with exactly one assigned gRNA (B), the median negative log2 fold change of the target gene (C), the number of significant target genes (D), the number of total discoveries on a data subset (E) and the number of false discoveries (F) is shown over various thresholds (x-axis). The black dotted lines mark the chosen threshold per method based on the maximum number of significant target genes.

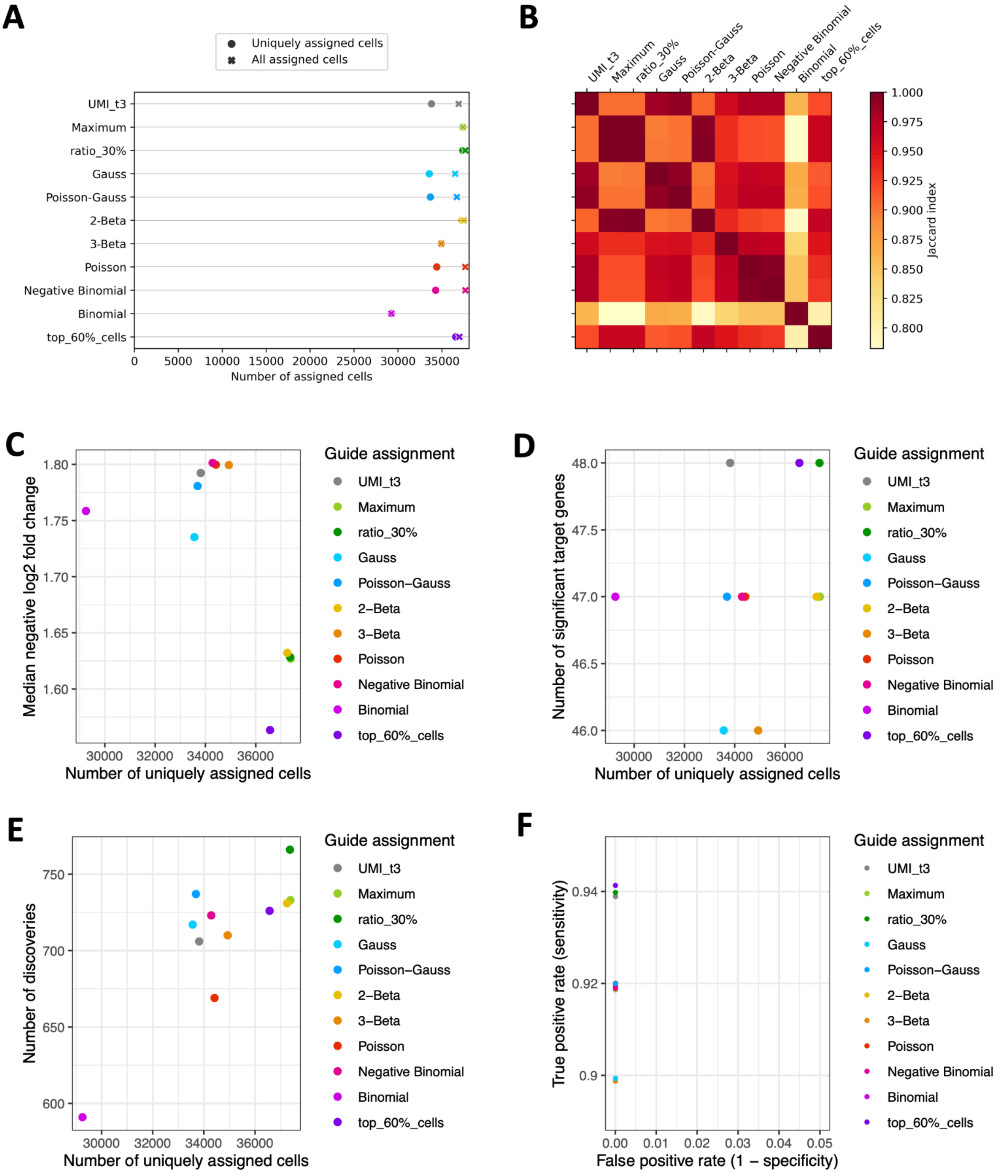


**Supplementary Figure 12: Method comparison for the whole transcriptomics screen by Schraivogel et al.** A: Number of total assigned cells (crosses) and cells with a single gRNA assigned (filled circle) per method. B: Assignment similarity heatmap showing the Jaccard index of the assignments per method pair. Only cells with one assigned gRNA were considered. C-E: Median negative log2 fold change of the target gene per gRNA (C), number of significant targets (D) and number of total discoveries (E) (y-axis) vs. the number of uniquely assigned cells per method (x-axis). The number of total discoveries was calculated for 5000 response genes (see Supplementary Information for details). F: True positive rate (y-axis) vs. false positive rate (x-axis) of all methods. The y-axis has been scaled discrete and the points were slightly shifted up and down to avoid overplotting.

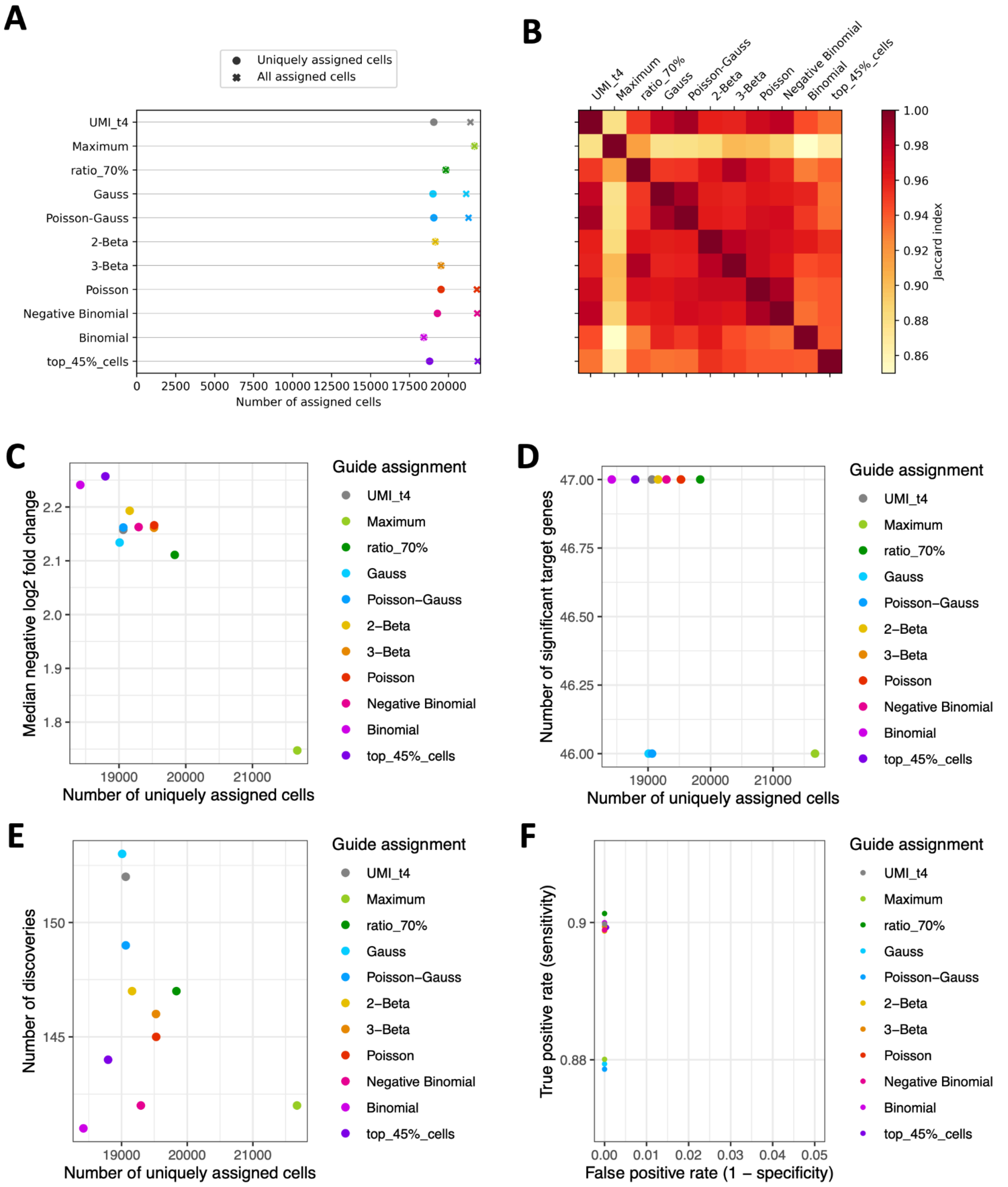


**Supplementary Figure 13: Method comparison on the TAPseq screen by Schraivogel et al.** A: Number of total assigned cells (crosses) and cells with a single gRNA assigned (filled circle) per method. B: Assignment similarity heatmap showing the Jaccard index of the assignments per method pair. Only cells with one assigned gRNA were considered. C-E: Median negative log2 fold change of the target gene per gRNA (C), number of significant targets (D) and number of total discoveries (E) (y-axis) vs. the number of uniquely assigned cells per method (x-axis). F: True positive rate (y-axis) vs. false positive rate (x-axis) of all methods. The y-axis has been scaled discrete and the points were slightly shifted up and down to avoid overplotting.
